## Supplementary for "The Surprising Role of the Default Mode Network"

**Supplementary Note 1**

As stated in the main text and demonstrated in Supplementary Figure 2, significant correlations (via significance test; $p < 0.05$) were found between surprise ratings and ISFC among DMN region pairs for Bang! You're Dead, similar to correlation pattern observed in Sherlock. However, unlike in Sherlock, in Bang! You're Dead, a similar pattern of significant correlations ($p < 0.05$) was found also for emotional intensity ratings. To test whether this was a result of similar ratings across the two behavioral measures throughout the movie, we correlated across the time-courses of surprise and emotional intensity, revealing a very high correlation ($R = 0.92$, $p < 0.001$). This suggests that surprise was strongly confounded with emotional intensity during Bang! You're Dead, which is a thriller movie. We thus did not extend the analysis of these data, as the cognitive states could not be disentangled.

| Measure | Questionnaire Instructions | Scale min – 1 | Scale max – 7 |
| --- | --- | --- | --- |
| **Vividness** | Please take a moment to recall this moment of the movie. How **vivid** is your memory of this event? | cloudy and imageless | clear and vivid as if experienced again |
| **Free recall** | Focus on this particular moment in the movie, including no more than a few seconds before and after the described event. Write down every detail you can remember, including any or all of the following types of information: what happened in the movie, what you saw, what you heard, what were your own thoughts, emotions and/or physical sensations while you were watching that event, etc. | n/a | n/a |
| **Surprise** | How **surprising** was the event? | did not surprise me at all | no other event in the movie surprised me this much |
| **Emotional intensity** | How **emotionally intense** was the event while watching it? | no detectable emotion | the most intense event to watch in this movie |
| **Emotional valence** | Would you rate this event as **emotionally positive or negative**? | strongly negative | strongly positive |
| **Importance** | How **important** was this event **to the main story** of the movie? | insignificant | more important than any other event in the movie |

**Supplementary Table 1. Questionnaire phrasing of scale rating and free recall instructions.** Instructions were repeated identically with every event reminder. Scale ratings were reported via keys 1 through 7 on the computer keyboard. Free recall was typed into an open-ended response field under a 3-minute time-limit per event reminder.

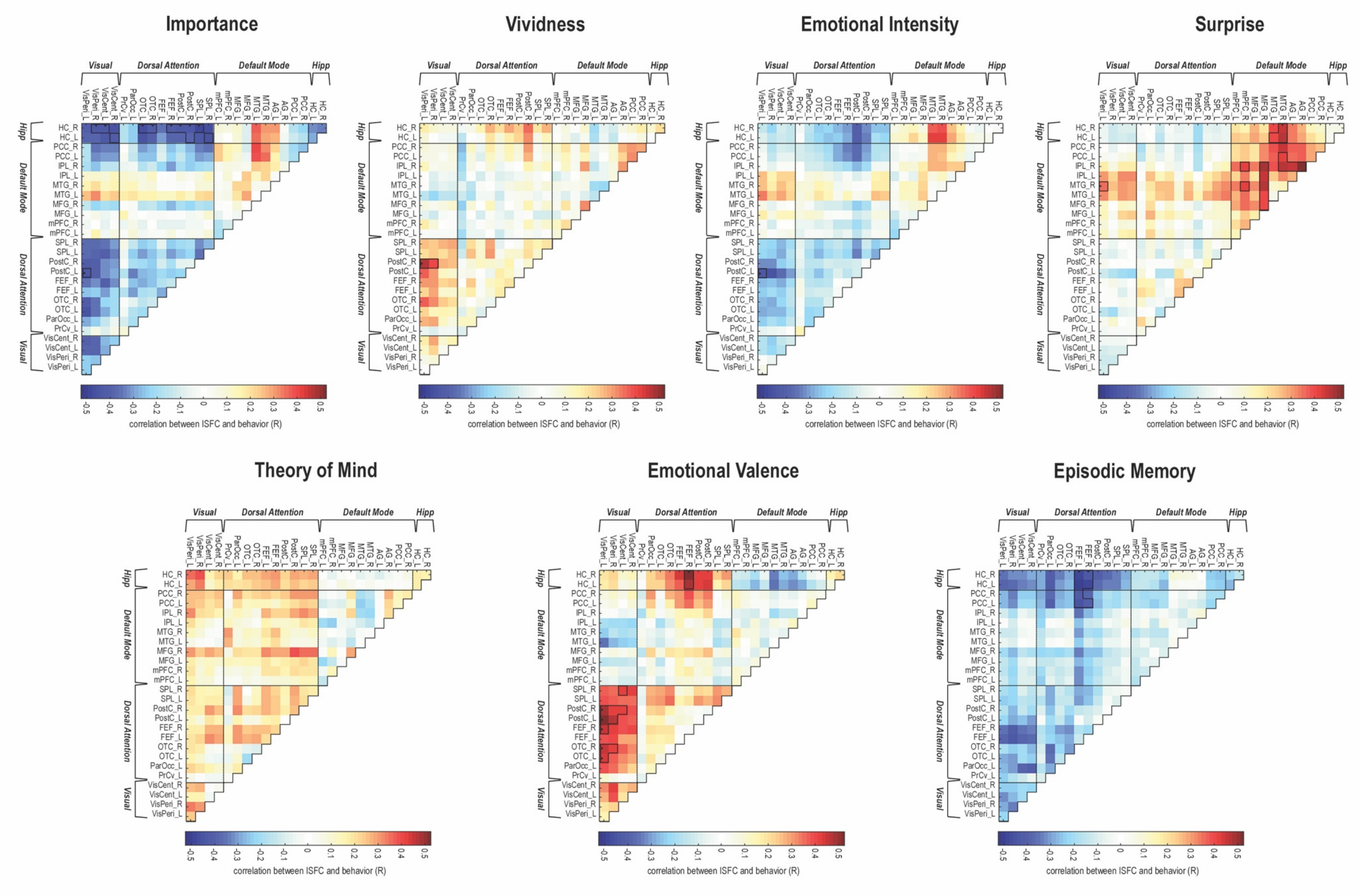
Importance
Vividness
Emotional Intensity
Surprise
Theory of Mind
Emotional Valence
Episodic Memory
Visual
Dorsal Attention
Default Mode
Hipp
correlation between ISFC and behavior (R)

**Supplementary Figure 1. Correlation between the fluctuation in each behavioral measure and fluctuation in coactivations of every region pair for Sherlock.** Correlation SFPA – pairwise ISFC time-courses (mean of 35 fMRI participants) were tested for correlation with the time-course of each of the 7 behavioral measures (mean of 45 behavioral participants). Black outlines denote above-chance correlations at $p < 0.05$ (corrected), determined by random permutation testing (1000 iterations).

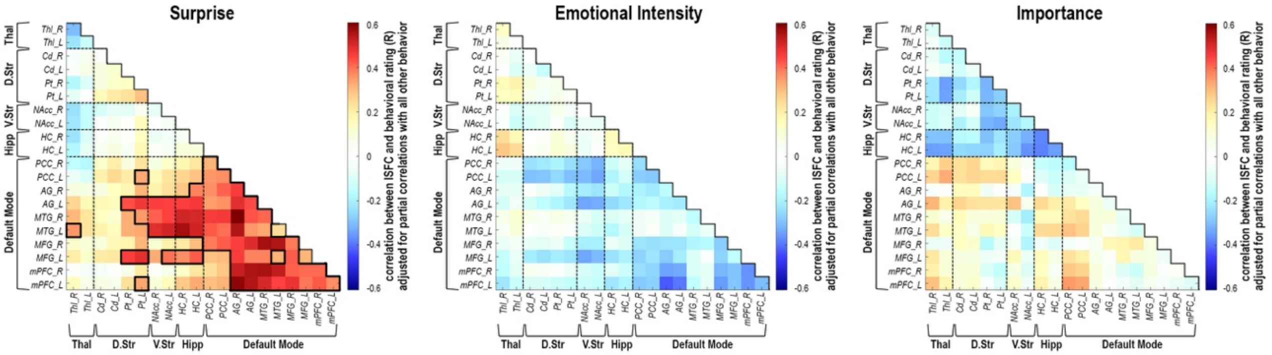


**Supplementary Figure 2. Correlation analysis adjusted for partial correlations among behavioral measures in Sherlock.** To assess the contributions of collinearities among behavioral measures to SFPA results, we performed correlation SFPA as in the original analysis, but adjusted for partial correlations between each behavioral measure and all others. Measures of highest behavioral collinearity with surprise were emotional intensity and importance. Results show that after adjusting for partial correlations among all behavioral measures, surprise is significantly associated with ISFC of the DMN and subcortical regions, whereas emotional intensity and importance show no above-chance SFPA. Pearson correlations were calculated between each behavioral time-course (mean of 45 behavioral participants), adjusted for partial correlations with all other measures, and ISFC of each region-pair (mean of 35 fMRI participants), across the time-course of movie events. Black outlines denote above-chance correlations at $p < 0.05$ (corrected), determined by random permutation testing (1000 iterations).

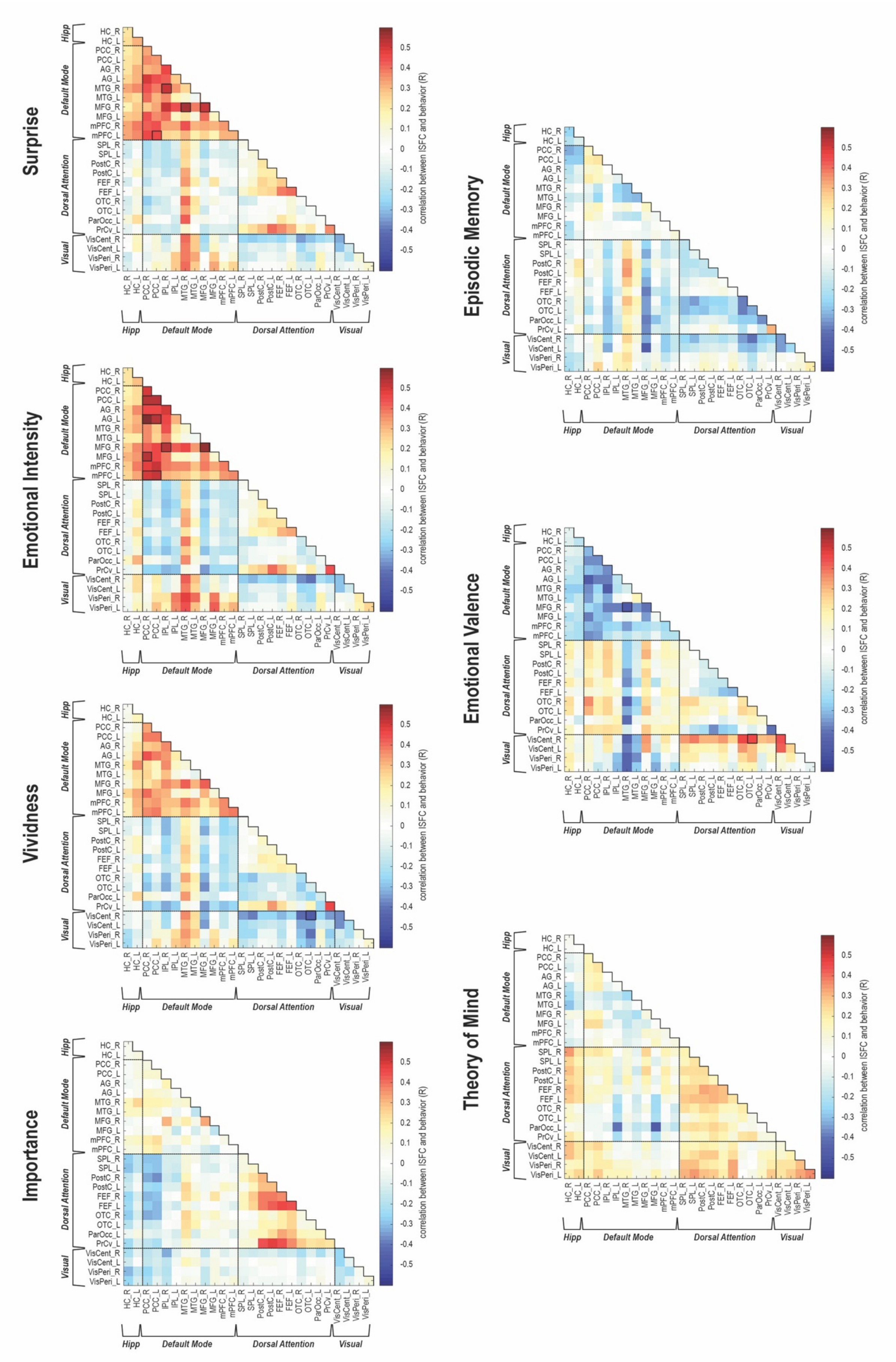
Surprise
Emotional Intensity
Vividness
Importance
Episodic Memory
Emotional Valence
Theory of Mind
Hipp
Default Mode
Dorsal Attention
Visual
correlation between ISFC and behavior (R)

**Supplementary Figure 3. Correlation between the fluctuation in each behavioral measure and fluctuation in coactivations of every region pair for Bang! You're Dead.** Correlation SFPA – pairwise ISFC time-courses (mean of 30 fMRI participants) were tested for correlation with the time-course of each of the 7 behavioral measures (mean of 42 behavioral participants). Black outlines denote above-chance correlations at $p < 0.05$ (corrected), determined by random permutation testing (1000 iterations).

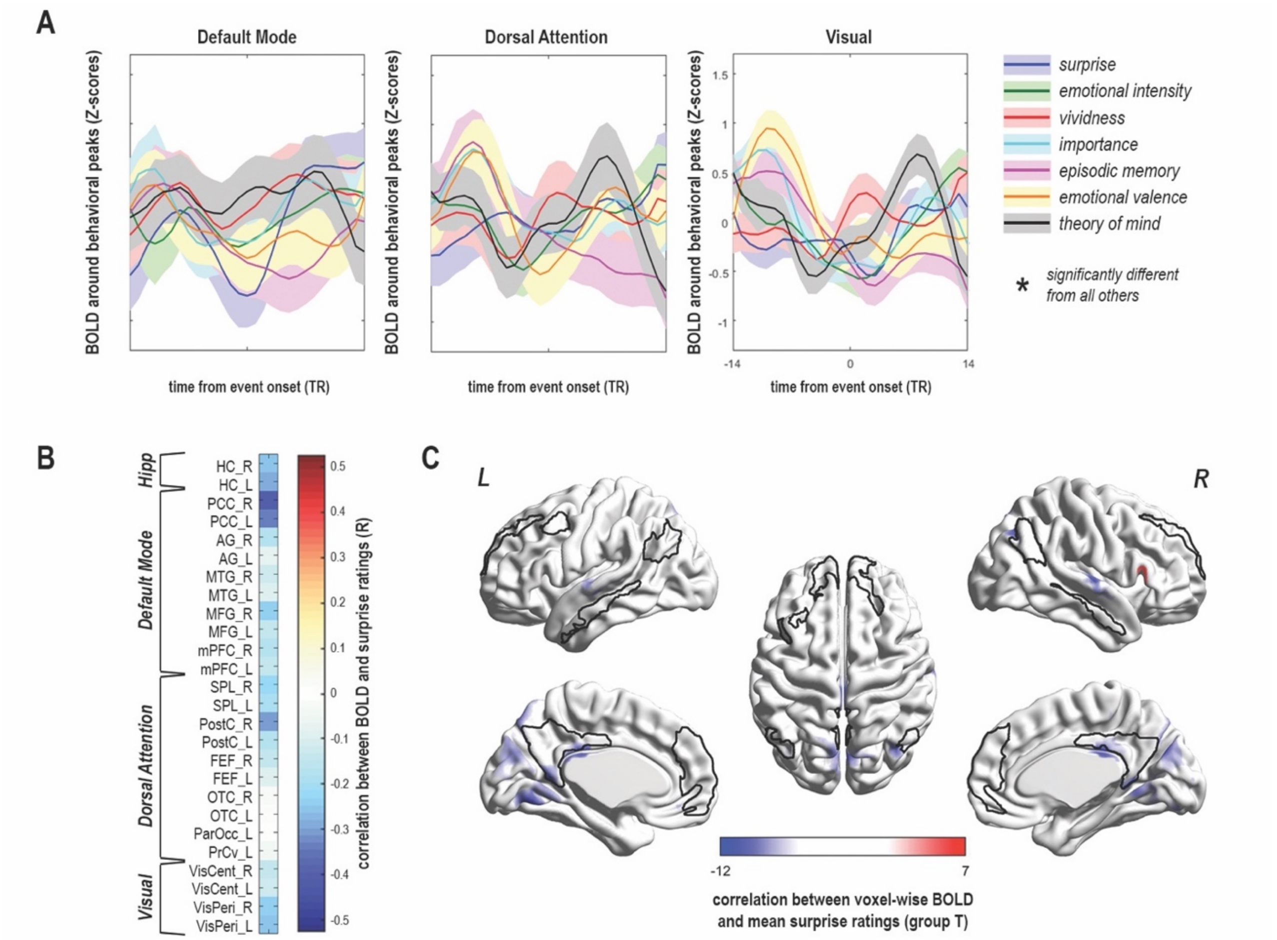


**Supplementary Figure 4. Univariate activations in Sherlock.** (A) Peak SFPA shows the univariate network BOLD averaged across the 5 peak events on each behavioral measure, in DMN (left), DAN (middle) and Vis (right). No significant differences in BOLD seen between different peak states, as examined by random permutation testing (1000 iterations) at $p < 0.05$. Network BOLD is plotted as mean ± SEM across subjects. Time 0 is the scan volume corresponding to the event, plotted ± 14 volumes adjacent to event. (B) Correlation SFPA, between the time-course of surprise ratings and the time-course of region-wise univariate BOLD. No above-chance correlations seen, as examined by random permutation testing (1000 iterations) at $p < 0.05$ (corrected). (C) Whole-brain correlations between the time-course of surprise ratings and the time-course of voxel-wise univariate BOLD. Above-chance negative correlations shown in blue, above-chance positive correlations shown in red, as tested across subjects by T-test of fisher-transformed correlation coefficients, at $p < 0.05$ (corrected). Black outlines denote the ROIs of the DMN, demonstrating little to no overlap with univariate effects of surprise. Univariate BOLD calculated across 35 fMRI participants; Behavioral ratings calculated across 45 behavioral participants.

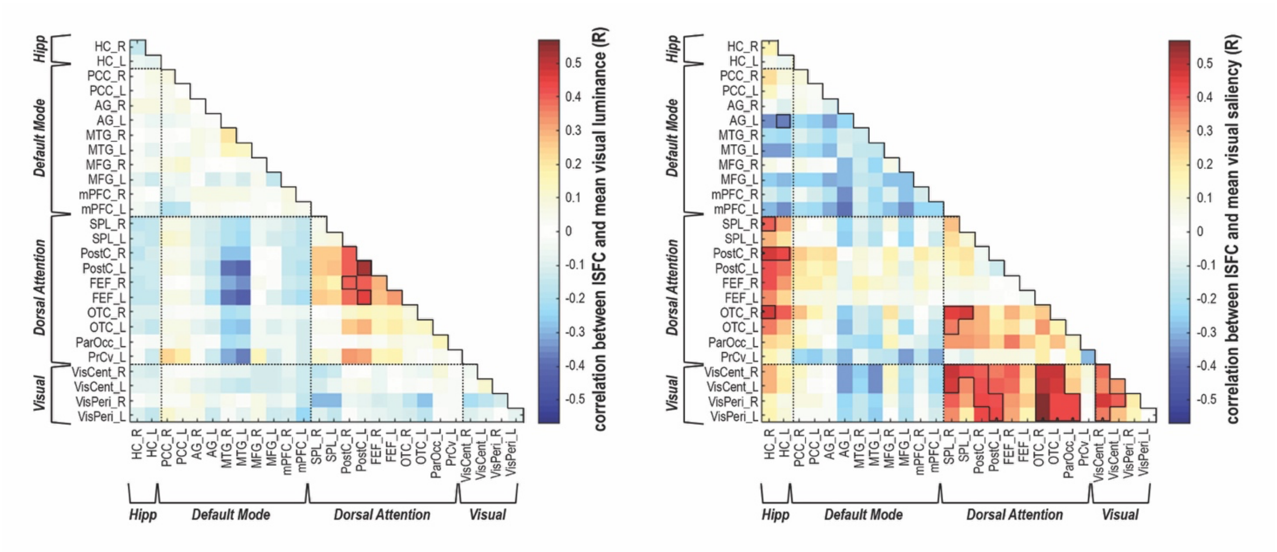


**Supplementary Figure 5. Correlation between fluctuation in visual attributes and fluctuation in coactivations of every region pair for Sherlock.** Fluctuations in mean visual luminance (left) and mean visual saliency (right), extracted from movie frames across each event window, were tested for correlation with the fluctuation in ISFC between each pair of regions (mean of 35 fMRI participants). Black outlines denote above-chance correlations at $p < 0.05$ (corrected), determined by random permutation testing (1000 iterations).

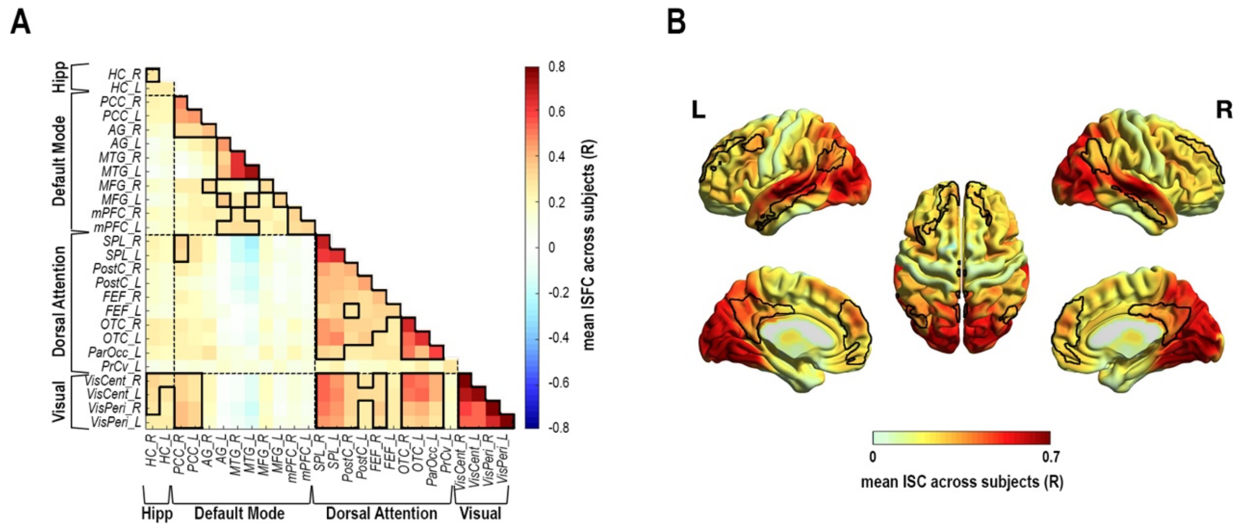


**Supplementary Figure 6. Inter-subject correlations across the entire scan time-course for Sherlock.** (A) ISFC between every pair of ROIs, calculated as the Pearson correlation across all 946 scanning volumes. Black outlines denote above-chance correlations at $p < 0.05$, determined by random permutation testing (1000 iterations); (B) Voxel-wise ISC throughout the whole brain, calculated as the Pearson correlation across all 946 scanning volumes. Black outlines denote the ROIs of the DMN. ISFC and ISC values plotted as means of 35 fMRI participants.

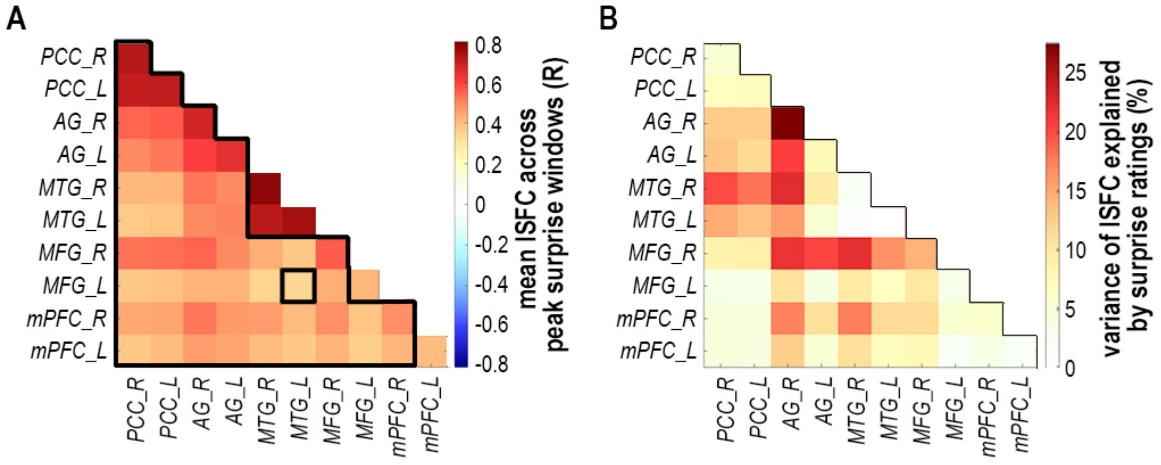


**Supplementary Figure 7. Inter-subject functional correlation (ISFC) at peak surprise and surprise-explained variance in the default mode network (DMN) for Sherlock.** (A) Peak-SFPA for each DMN region-pair, as the ISFC values (mean of 35 fMRI participants) at event onset, averaged across the 5 peak surprising events. Black outlines denote above-chance SFPA ($p < 0.05$) via permutation testing (1000 iterations); (B) The percentage of variance explained by surprise ratings in ISFC fluctuation, for each DMN region pair. Explained variance is calculated as the squared Pearson correlation between surprise ratings (mean of 45 behavioral participants) and ISFC (mean of 35 fMRI participants) across the 49 movie events, plotted as percentage ($R^2$ x 100).

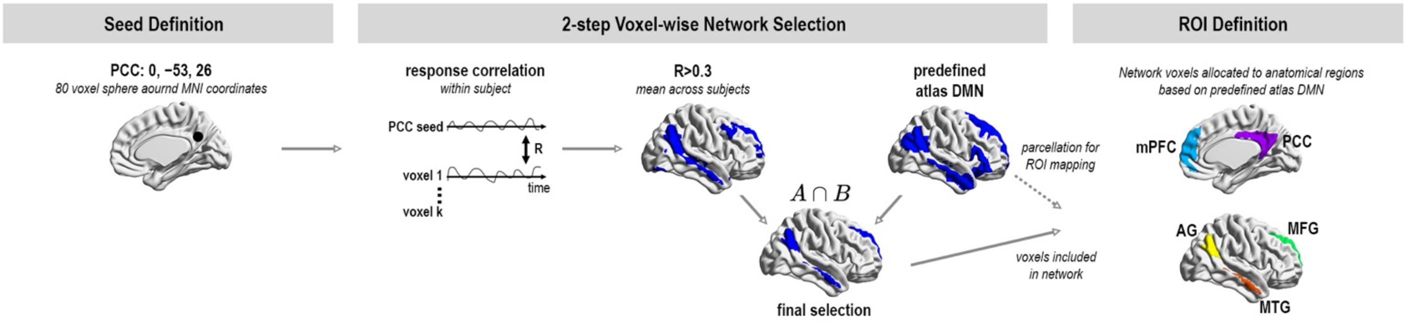


**Supplementary Figure 8. Region of interest (ROI) localization.** Workflow, from left to right: 1. Seed for functional localization of the network was defined anatomically as a sphere around MNI coordinates validated in previous reports[1]; 2. Whole-brain voxel-wise network definition, first, precluded voxels of low mean response correlation with the seed region (during a non-target scan), and second, precluded voxels mapped outside a network atlas that had been functionally defined based on a wide sample[1]. Voxels passing both filtering stages were selected for the final network definition; 3. For ROI definition, network voxels were allocated to general anatomical regions mapped to the predefined network atlas[1].

[1]Schaefer, A. et al. Local-Global Parcellation of the Human Cerebral Cortex from Intrinsic Functional Connectivity MRI. Cereb Cortex 28, 3095-3114, doi:10.1093/cercor/bhx179 (2018).

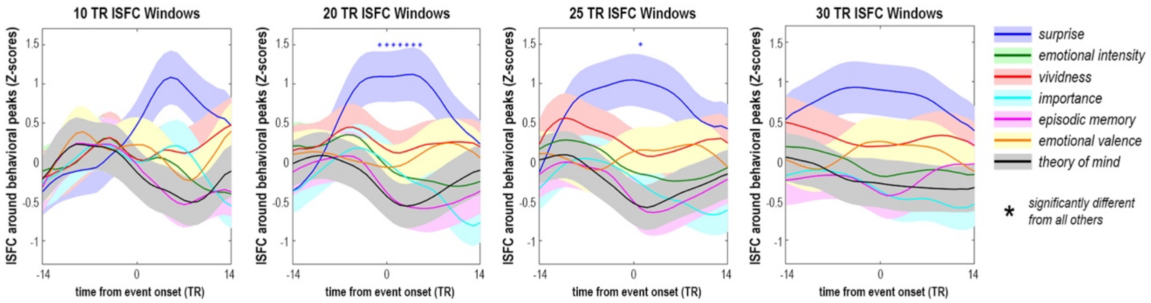


**Supplementary Figure 9. DMN peak analysis for Sherlock using different time-window sizes.** ISFC was calculated across 10, 20, 25, or 30 TR windows. Peak SFPA – ISFC mean of 35 fMRI participants and of all network regions was averaged across the 5 peak events on each behavioral measure (e.g. ISFC during 5 most surprising events). This resulted in a mean ISFC value per network per peak-state, presented here. DMN regions were selectively coactivated during peak surprise, compared to all other peak states, as revealed by random permutation testing (1000 iterations) at $p < 0.05$. Network ISFC is plotted as mean ± SEM across subjects, corresponding to event onset (time 0), and to each of the 14 TRs before and after the event, during each peak state (irrespective of window size for ISFC calculation, described above).
